# A versatile two-step CRISPR- and RMCE-based strategy for efficient genome engineering in *Drosophila*

**DOI:** 10.1101/007864

**Authors:** Xu Zhang, Wouter Koolhaas, Frank Schnorrer

**Affiliations:** Max Planck Institute of Biochemistry, Am Klopferspitz 18, 82152 Martinsried, Germany

## Abstract

The development of CRISPR/Cas9 technologies promises a quantum leap in genome-engineering of model organisms. However, CRISPR-mediated gene targeting reports in *Drosophila* are still restricted to a few genes, use variable experimental conditions and vary in efficiency, questioning the universal applicability of the method. Here, we developed an efficient, two-step strategy to flexibly engineer the fly genome by combining CRISPR with recombinase-mediated cassette exchange (RMCE). In the first step, two sgRNAs, whose activity had been tested in cell culture, were co-injected together with a donor plasmid into transgenic *Act5C-Cas9, Ligase4* mutant embryos and the homologous integration events were identified by eye fluorescence. In the second step, the eye marker was replaced with DNA sequences of choice using RMCE enabling flexible gene modification. We applied this strategy to engineer four different loci, including a gene on the fourth chromosome, at comparably high efficiencies, suggesting that any fly lab can engineer their favourite gene for a broad range of applications within about three months.

---

Reverse genetics is currently booming with the establishment of TALEN-and CRISPR-mediated genome engineering^1–3^. In particular, the CRISPR/Cas9 technology appears to efficiently and specifically introduce double strand DNA breaks in the genome of the organism, which can then be utilised to either introduce point mutations by error prone nonhomologous end-joining (NHEJ) or to integrate heterologous DNA into the chromosome using the homology-directed repair (HDR) pathway^2^. In *Drosophila*, CRISPR-induced NHEJ technology has mainly been utilised to mutate genes that result in a visible, easily scored phenotype, such as white eyes or yellow body colour, or to mutate GFP transgenes^4–7^. Mutants in genes with no visible phenotype required laborious PCR screening for their identification^8–10^, thus far hampering CRISPR-based gene mutagenesis to develop into a routine application. Recently, this bottleneck was addressed by applying CRISPR-induced HDR to insert an attP-site together with a visible marker into the gene of interest^11–13^. In some cases the visible marker was flanked by FRT or loxP sites allowing its excision to only leave one attP site (and one loxP or FRT site) within the gene. This attP site enables the introduction of any given DNA sequence into the gene of interest^11, 12^. However, the efficiency of reporter integration was rather low^11^ and only determined at a single genomic locus^12, 13^, leaving the general applicability to the *Drosophila* genome an open question.

Here, we have developed a highly flexible two-step genome engineering platform that combines CRISPR-mediated HDR with ΦC31 recombinase-mediated cassette exchange (RMCE). In the first step, CRISPR is applied to integrate a splice acceptor and an SV40 terminator together with a 3xP3-dsRed eye reporter. This enables both, the efficient identification of the targeted event and the creation of a strong loss of function allele. In the second step, two flanking attP sites are utilised to replace the inserted DNA by any DNA of choice using RMCE, an established standard technology in *Drosophila*^14^. Together, this allows flexible cassette exchange to freely manipulate the gene of interest. We successfully applied this method to four different loci and efficiently generated several allele variants, including a conditional allele, from a single targeting event. Our streamlined CRISPR/Cas9-and RMCE-based strategy makes it practical to flexibly engineer any *Drosophila* gene of choice for a broad range of applications within about three months.

## Results

### Strategy overview

We aimed to develop a versatile and efficient strategy to modify the *Drosophila* genome, which would allow various genome modifications such as the introduction of single point mutations, protein tags, exon deletions or other desired changes in the gene of choice. Despite the suggested higher efficiency of CRISPR/Cas9-induced HDR as compared to Zn-finger-or TALEN-induced HDR, the identification of successfully targeted carrier flies is still a limiting step in the process. PCR-based screening or melting curve analysis methods require DNA extraction^15^ and thus are laborious and often inconvenient for efficient stock generation. Therefore, we decided to develop a 2-step strategy as illustrated in Figure 1, which enables efficient identification of the carrier flies and allows entirely flexible genome engineering. In the first step, we insert a 3xP3-dsRed marker enabling easy identification of the HDR event. A strong splice acceptor, followed by STOP codons and an SV40 polyA terminator, precedes the dsRed cassette. The inserted DNA is flanked by two attP sites in opposite orientations, a strategy that we adapted from the popular MiMIC system^14^. If this cassette is inserted into an intron, or replaces an endogenous exon, it results in a truncated mRNA of the targeted gene. Thus, step 1 creates a loss of function allele (Figure 1).

**Figure 1.**
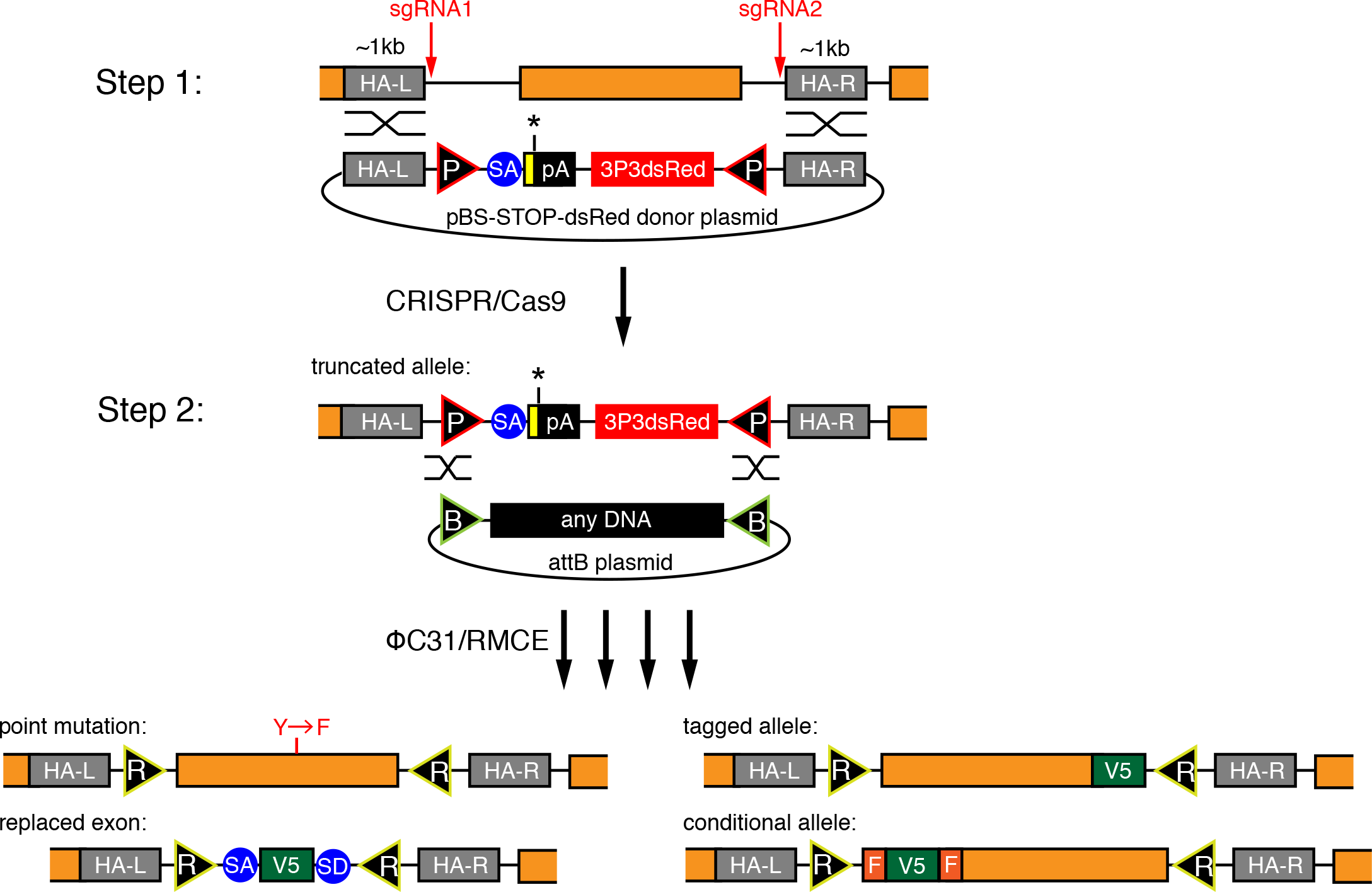
| A 2-step method to flexibly engineer the fly genome. Overview of the 2-step procedure. In step 1, a donor vector consisting of two attP sites (P), a splice acceptor (SA), STOP codons (yellow box, black asterisk) followed by an SV40 polyadenylation signal (pA) and a 3xP3dsRed marker are inserted upon Cas9 cleavage with two sgRNAs. The orange coding exon is excised. In step 2, ΦC31-mediated RMCE inserts any DNA sequence between the 2 attB sites (B). Examples for various engineered exons are given, resulting in attR sites (R) in introns.

In the second step, RMCE is applied to replace the DNA between both attP sites by any DNA of choice, leaving a minimal scar of two attR sites, preferably in introns. RMCE has been used very efficiently in the MiMIC system, demonstrating that attR sites in introns generally do not interfere with gene function^14^. Hence, our strategy enables the generation of various alleles, like a defined point mutation, a tagged allele, an exon replaced by a tag or a conditional allele, from a single HDR carrier (Figure 1). This strategy should allow flexible editing of any *Drosophila* gene within about three months (Figure 2).

**Figure 2.**
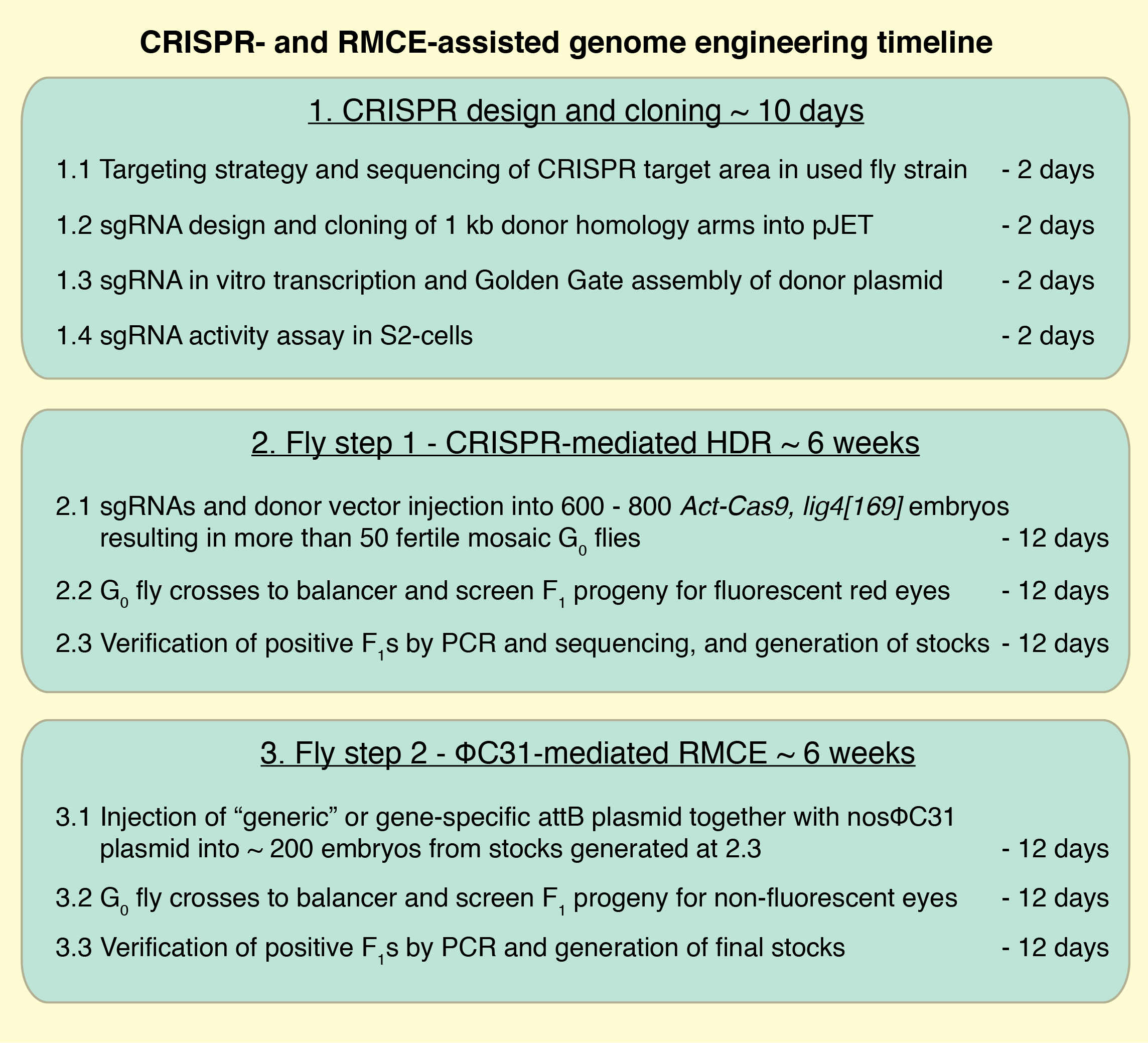
| 2-step genome engineering timeline. Schematic overview of the major steps of the genome engineering procedure. Details are provided in the Methods section.

### CRISPR design and cloning

In step 1 we aimed to insert a STOP-3xP3-dsRed cassette flanked by two attP sites using a donor plasmid (Figure 1). Since the same strategy should be applicable to any gene, we established a single step Golden Gate protocol to assemble the STOP-dsRed donor plasmid containing about 1kb homology arms on each side, which has been shown to be of sufficient length for efficient HDR^16^ and can be easily amplified by PCR. Cloning of the homology arms into the donor vector is thus very straightforward and takes only a few days for the gene of choice (Figure 3a, see Methods). This donor vector is the template for the HDR in step 1.

**Figure 3.**
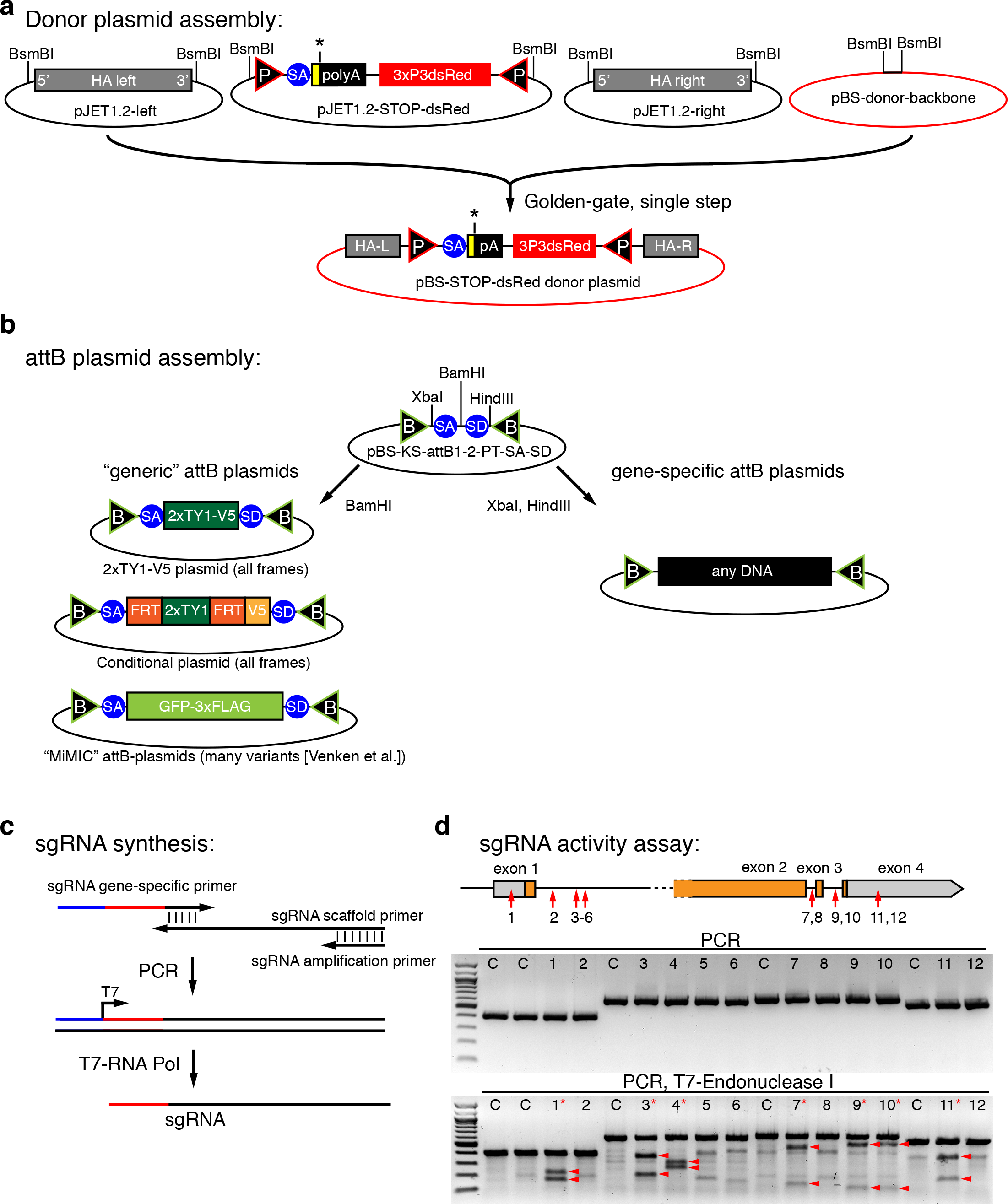
| Cloning scheme and sgRNA activity tests. (**a**) Single step golden gate assembly scheme of the STOP-dsRed donor vector cloned into a modified pBluescript backbone. (**b**) Scheme of the “generic” attB plasmids used in our study. A simple cloning step is sufficient to generate any gene-specific attB plasmids that can be used to replace an excised exon. (**c**) sgRNA synthesis scheme. (**d**) sgRNA activity assay of 12 different sgRNAs in S2 cells. PCR result with (bottom) or without (middle) T7-Endonuclease I treatment are shown. Digested products are marked by arrow heads and effective sgRNAs are marked by a red asterisk.

In step 2 RMCE exchanges the STOP-dsRed by any sequence located between 2 attB-sites in a provide plasmid (Figure 1). RMCE works very reliably and a large collection of plasmids to tag genes or insert reporters for various applications is available^14^. These plasmids are fully compatible with our step 2 design. We have generated additional “generic” attB plasmids that can be used to tag any gene with a 2xTY1-V5 tag or to engineer a conditional allele using an FRT flanked 2xTY1 cassette followed by a V5 tag (Figure 3b). Flp mediated deletion of the 2xTY1 cassette will lead to a frame shift and can thus be used to create loss of function clones at very high efficiency, as the flip-out will occur in cis^17^. We have generated both constructs in all three reading frames.

### CRISPR activity assay in cell cuture

Many search algorithms exist to predict sgRNA target sequences for a given gene region^18^. However, to date there is no simple way of confirming if any of the predicted sgRNAs work efficiently. We developed such a selection assay to be able to only inject effective sgRNAs into fly embryos. We designed 12 different sgRNAs targeting different regions in the *salm* gene and synthesized the sgRNAs by a standard PCR and in vitro transcription reaction (Figure 3c). These sgRNAs were then individually transfected into Cas9 expressing S2 cells^19^, and their cleavage efficiency was determined with a simple T7-Endonuclease I assay (see Methods). On average, about half of the tested sgRNAs work efficiently in this assay (Figure 3d and data not shown), strongly suggesting that such a pre-selection test is useful to improve the *in vivo* success rates.

### Step 1 - HDR in *Lig4* mutant embryos

To test the efficiency of inserting our STOP-dsRed cassette, we designed three donor constructs targeting different regions in the *salm* gene (Figure 4a): the first, deleting parts of exon 1; the second, inserting the cassette into intron 1; and the third, deleting exon 3. For each construct about 1 kb homology arms were cloned in the STOP-dsRed donor vector. We injected the STOP-dsRed donor as circular plasmid together with 2 plasmids each containing a U6 promoter driven sgRNA verified in S2 cells and a hsp70-Cas9 source (see Methods). We injected into *Ligase4* mutant embryos, which were reported to exhibit a higher rate of HDR than wild-type embryos^15, 16^. We injected between 700 and 1500 embryos for each of the three constructs and were able to recover 11 red eyed F1 carriers from 2 independent founders for the first intron construct and 72 red eyed F1s from 4 independent founders for the third exon deletion construct (Table 1). This demonstrated that our strategy works in principle, but as we failed to recover the first exon deletion allele, we wanted to further improve the efficiency by using a different Cas9 source.

**Figure 4.**
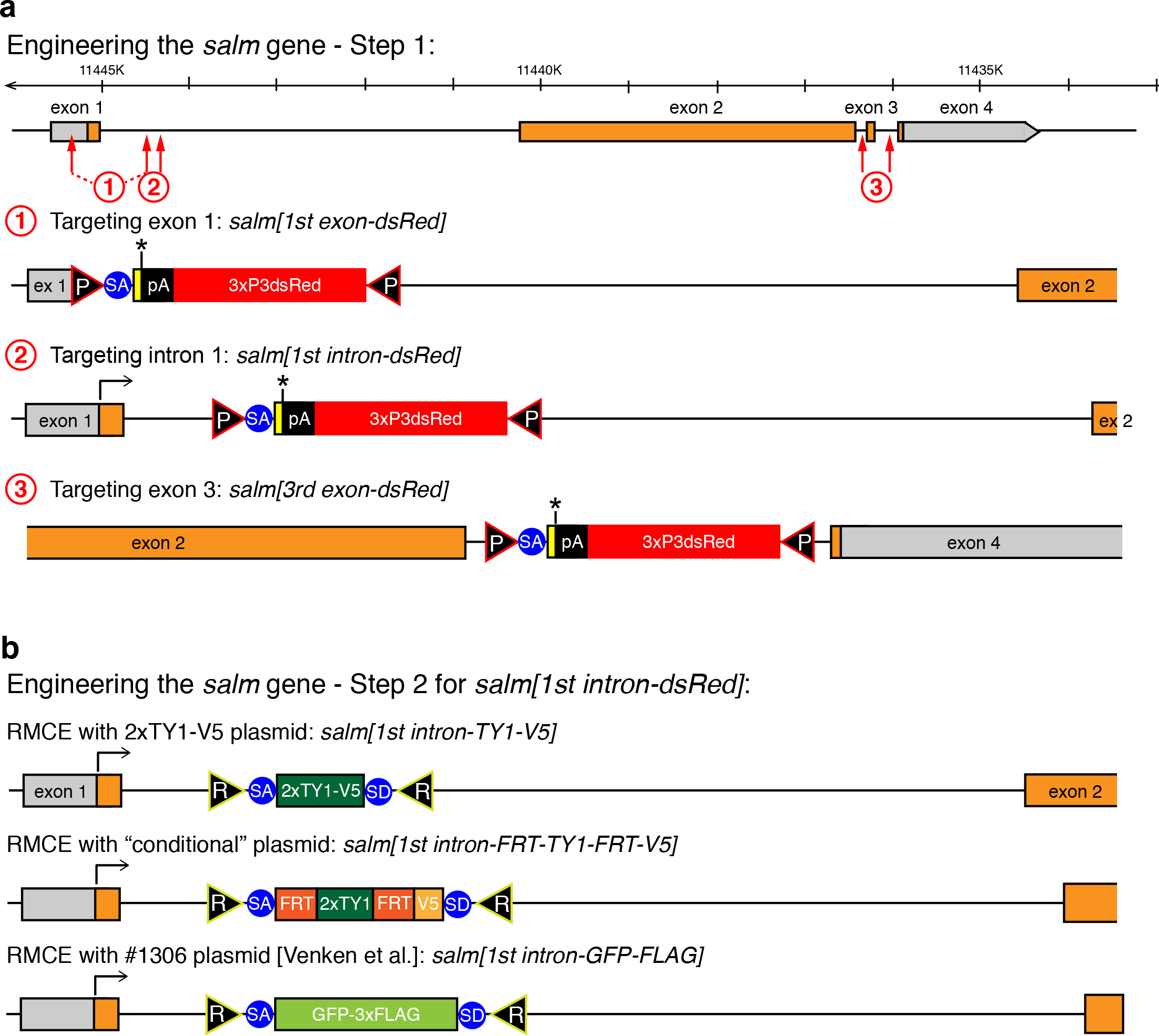
| Engineering of the *salm* gene. (**a**) Step 1 engineering of the *salm* gene. The genomic *salm* organisation is depicted with coding exons in orange. The sgRNA targeting sites are indicated by red arrows and the resulting *salm[1^st^ exon-dsRed], [1^st^ intron-dsRed]* and *[3^rd^ exon-dsRed]* alleles are shown. (**b**) Step 2 engineering of the *salm* gene. RMCE products of the *salm[1^st^ intron-dsRed]* with three different exon cassettes are shown.

**Table 1.** | Summary of the transformation efficiencies for the four different loci modified in this study.

| Locus | Injected genotype | Injected embryos | Larvae | G0 adults | Fertile G0 adults | Independent F1 carriers | Total red eyed F1 | Independent carriers per fertile G0s |
| --- | --- | --- | --- | --- | --- | --- | --- | --- |
| *salm* 1st exon | *Lig4* -/- | 1500 | 396 | 166 | n.d. | **0** | 0 | n.d. |
| *salm* 1st intron | *Lig4* -/- | 700 | 171 | 149 | n.d. | **2** | 11 | n.d. |
| *salm* 3rd exon | *Lig4* -/- | 1500 | 312 | 150 | n.d. | **4** | 72 | n.d. |
| *salm* 1st exon | *Act5C-Cas9, Lig4* -/- | 700 | 124 | 48 | 30 | **1** | 13 | **3,3%** |
| *salm* 1st intron | *Act5C-Cas9, Lig4* -/- | 700 | 200 | 122 | 64 | **9** | 54 | **14,1%** |
| *salm* 3rd exon | *Act5C-Cas9, Lig4* -/- | 700 | 291 | 150 | 99 | **4** | 59 | **4,0%** |
| *bent* 11th exon | *Act5C-Cas9, Lig4* -/- | 700 | 204 | 118 | 56 | **1** | 5 | **1,8%** |

### Step 1 - transgenic Cas9 improves HDR efficiency

A number of transgenic Cas9 flies have been recently generated, some of which have been used successfully^6, 12, 13^. To test weather a transgenic Cas9 source is indeed more efficient for HDR than a source from an injected plasmid, we targeted the same loci as above, but now using *Act5C-Cas9, Lig4* flies. For this, we recombined an *Act5C-Cas9* transgene expressing Cas9 ubiquitously, including maternally in the germ-line, with the *Lig4[169]* null allele. Additionally, we removed the *white* and *3xP3dsRed* markers from the *Act5C-Cas9* transgene to obtain a *Act5C-Cas9, Lig4[169]* chromosome that is useful for the injection of our donor plasmids (see Methods). We injected 700 *Act5C-Cas9, Lig4[169]* embryos with two *in vitro* transcribed sgRNAs targeting either the 1^st^ exon, the 1^st^ intron or the 3^rd^ exon of *salm* (using the same sgRNA target sequences as used above). We obtained 1, 9 and 4 independent founders producing 13, 54 and 59 F1 carriers, respectively, demonstrating that all 3 loci were targeted successfully with frequencies between 3 and 14 % per fertile G_0_ (Table 1). To verify that the targeted insertion occurred correctly, we tested a total of 9 independent carriers from the 3 loci by PCR and sequencing and were able to confirm all of them.

The first step of our gene targeting strategy inserts a strong splice acceptor followed by a STOP cassette into the gene and thus should terminate transcription at this position. By design, the *salm[1^st^ exon-dsRed]* allele additionally has a deleted ATG. As expected, the *salm[1^st^ exon-dsRed]* allele is homozygous lethal, as well as lethal in trans to *salm[1]*, demonstrating that we indeed created a strong *salm* loss of function allele (Figure 5a). The *salm[1^st^ intron-dsRed]* allele harbours an insertion in the first intron (Figure 4a). This allele is also homozygous lethal, and lethal in trans to *salm[1]* suggesting that the splice acceptor and STOP cassette are used efficiently to create a strong loss of function allele (Figure 5a). The *salm[3^rd^ exon-dsRed]* allele only deletes the last 36 amino acids of the long SalmPA isoform, including 10 amino acids of the last zinc finger (Figure 4a). This allele is homozygous viable, but flightless (Figure 5a), implying a role for the last zinc finger in flight muscle function. Taken together, these data suggest that our CRISPR-mediated step 1 strategy works efficiently to isolate targeted carrier flies at a practical frequency for routine use. Conveniently, these step 1 alleles are loss function alleles if the insertion is located within the gene.

**Figure 5.**
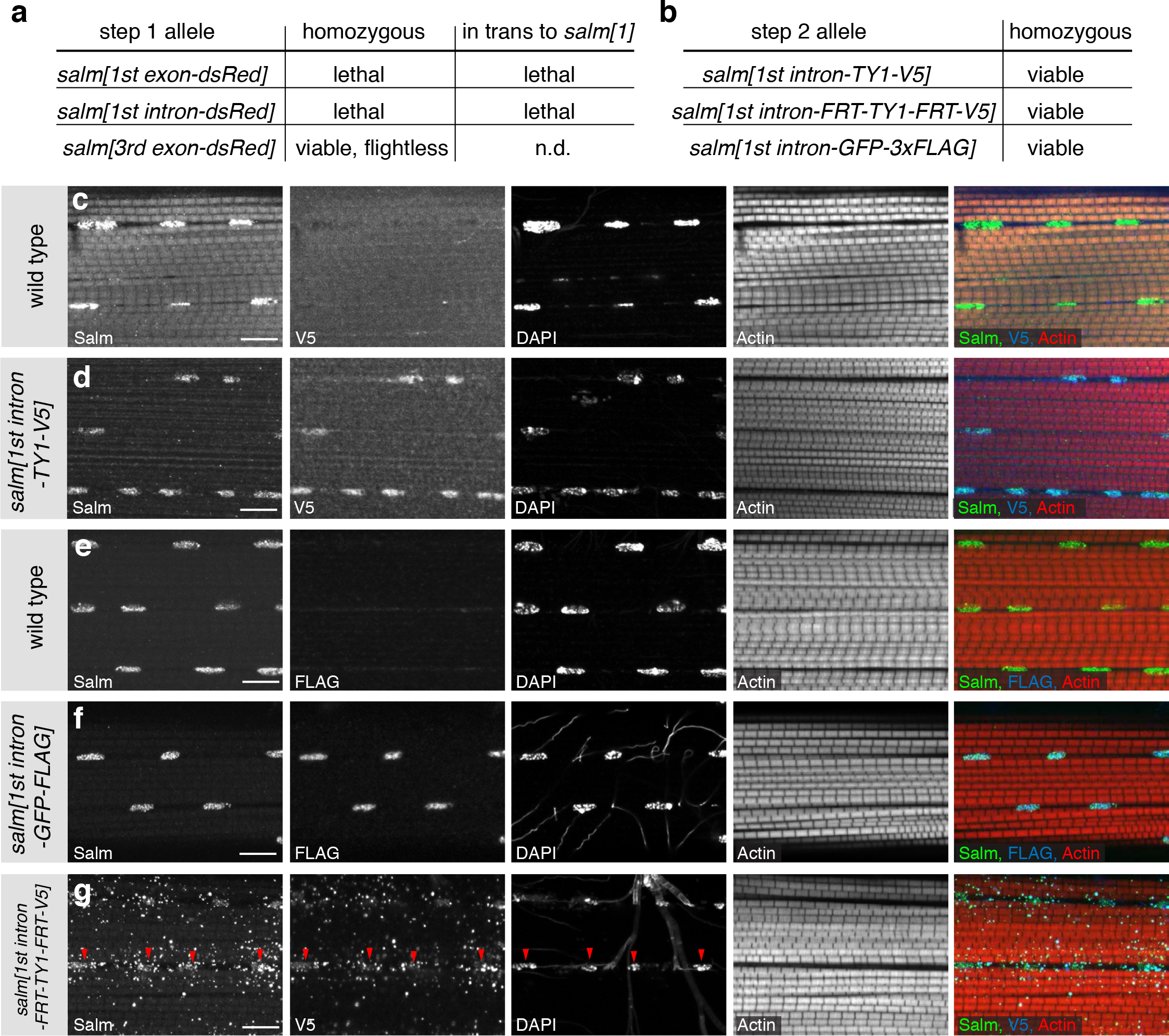
| Phenotypic analysis of the engineered *salm* alleles. (a) Lethality assay of the *salm-dsRed* alleles as homozygous or in trans to *salm[1]*. (b) All tagged *salm[1^st^ intron]* alleles regain homozygous viability after step 2. (**c-g**) Localisation of the tagged Salm proteins. Untagged Salm is located in the nucleus of wild type IFMs (c, e), whereas V5 tagged Salm is only detected in the *salm[1^st^ intron-TY1-V5]* and the *salm[1^st^ intron-FRT-TY1-FRT-V5]* alleles (d, g). FLAG is found in the IFM nuclei of *salm[1^st^ intron-GFP-FLAG]* adults (f). The Salm-FRT-TY1-FRT-V5 protein is found in the nuclei (red arrow heads) and also located in dots in the cytoplasm (g). Note the normal fibrillar morphology of the myofibrils in all the homozygous *salm[1^st^ intron]* alleles (d, f, g). Actin was stained with phalloidin and the scale bars are 10 µm.

**Figure 6.**
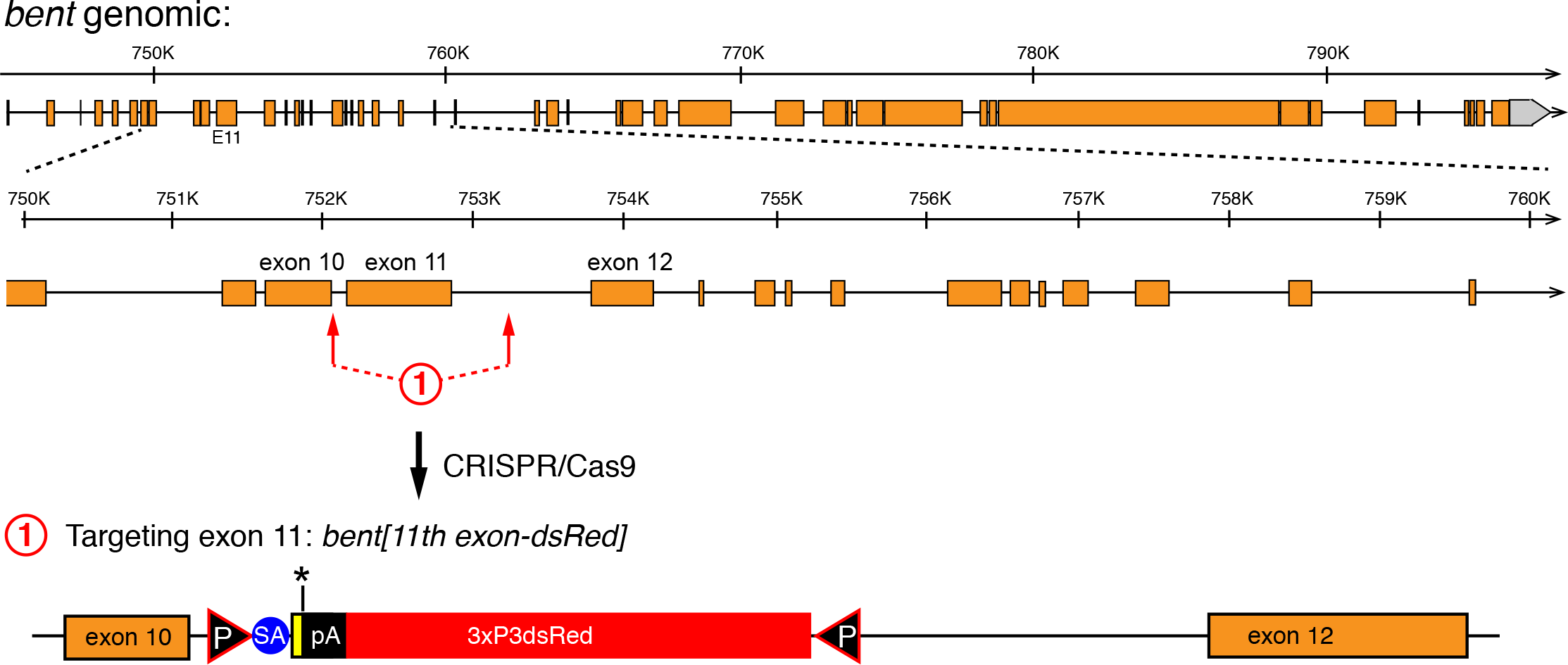
| Engineering of the *bent* gene. The entire genomic *bent* organisation is shown at the top with a 10 kb zoom-in below. Coding exons are in orange. The sgRNA targeting sites flanking exon 11 are indicated by red arrows and the resulting *bent[11^th^ exon-dsRed]* allele is shown at the bottom.

### Step 2 - Flexible gene editing by RMCE

A major benefit of our editing strategy is the flexible step 2 that enables the near-seamless insertion of any DNA sequence with only two remaining attR sites (Figure 1). To test the feasibility of step 2, we chose the *salm[1^st^ intron-dsRed]* allele. We exchanged the STOP-dsRed cassette with a short 2xTY1-V5 exon, a FRT-2xTY1-FRT-V5 conditional exon and a large GFP-3xFLAG exon from^14^ (Figure 4b). As expected, in all three cases the cassette exchange worked routinely (see Methods). Importantly, the *salm[1^st^ intron-dsRed]* lethality was reverted by RMCE in all three cases (Figure 5b). This demonstrates that our editing protocol generally does not result in any unwanted harmful mutations on the edited chromosome.

Salm protein is expressed in indirect flight muscles (IFMs) and is essential for fibrillar IFM fate specification^20^. Thus, we should detect the tagged Salm protein versions in the IFM nuclei of adult flies. Indeed, tagged protein from all three alleles, *salm[1^st^ intron-TY1-V5]*, *salm[1^st^ intron-FRT-TY1-FRT-V5]* and *salm[1^st^ intron-GFP-FLAG]* is expressed in IFMs. Salm-TY1-V5 and Salm-GFP-FLAG are readily detected in the IFM nuclei, Salm-FRT-TY1-FRT-V5 shows an additional dotty pattern in the cytosol (Figure 4c - g). The fibrillar IFMs organisation in all three homozygous *salm* alleles is normal showing that the tagged Salm proteins are indeed functional. Each IFM fiber contain several hundred nuclei. The conditional *salm[1^st^ intron-FRT-TY1-FRT-V5]* should now enable a clonal loss of function analysis of *salm* in muscle only, as flip out in cis is highly efficient^17^. Thus, this strategy should generally be versatile for the genetic analysis of muscle in the future.

### Gene editing on the fourth chromosome

In order to demonstrate the general applicability of our gene editing strategy, we decided to apply it to an additional locus. We chose the *bent* gene, located on chromosome four, which is highly heterochromatic and thus difficult to manipulate by standard genetic tools. To our knowledge, there is only a single case reported in the literature that targeted a gene located on the fourth chromosome by classical ends-out mediated homologous recombination using long homology arms^21^. *bent* encodes for a very large gene composed of at least 46 exons that are spread across more than 51 kb of genomic DNA (Figure 5). *bent* encodes for Projectin, a titin-like protein, which is specifically expressed in muscles and essential for correct sarcomeric organisation^22–24^. It is supposedly silent in germ cells in which the targeting event must happen. We chose to delete exon 11, an exon at the beginning of the PEVK domain of Projectin^25^, using two sgRNAs flanking the exon. Both sgRNAs tested positively in the S2 cell assay (Figure 5 and data not shown). We again used approximately 1kb homology arms and injected the donor vector into 700 *Act5C-Cas9, Lig4[169]* embryos. We isolated 5 carriers from 1 founder out of a total of 56 fertile G_0_ flies, an HDR efficiency of 1.8 %. We confirmed the *bt[11^th^ intron-dsRed]* allele by sequencing the locus. As expected, *bt[11^th^ intron-dsRed]* is homozygous lethal and also lethal in trans to *bt[I-b]*, a strong bent allele^23^, again suggesting that the inserted splice acceptor is used effectively and transcription is prematurely terminated. Together, these results demonstrate that our CRISPR-mediated targeting strategy also works efficiently on the fourth chromosome, suggesting it can be generally applied to any locus of choice in the fly genome.

## Discussion

CRISPR/Cas9 has been used successfully in many model organisms to generate mutants or to introduce targeted changes by HDR^1^. In *Drosophila*, there has been no general agreement regarding which strategy works most effectively to engineer the genome. To simply mutate a gene by CRISPR/Cas9-induced NHEJ, Cas9 was either injected as mRNA^7, 9^, provided from an injected plasmid^4, 11^ or provided from a transgenic source^5, 6, 8^. Similarly, the sgRNA was either injected as *in vitro* transcribed sgRNA, or provided by an injected plasmid or a transgenic source. A standard protocol has not yet emerged although a number of genes have been mutated.

NHEJ can only induce small insertions or deletions. In contrast, HDR allows the defined engineering of a given gene and thus is suitable for a much wider range of applications. CRISPR/Cas9-mediated HDR has been used in *Drosophila* to insert short attP or tag sequences from single strand oligonucleotides as donors^4^ or larger cassettes including a dsRed marker cassette from a plasmid donor^10–13^ again using various ways of injected or transgenic sources of sgRNAs or Cas9. The injected genotype was variable, sometimes *Lig4* mutants were used^10, 12^, sometimes not^4, 11–13^. Often the detection of the targeted event required laborious fly screening by PCR^10, 26^.

Here we aimed to develop a universal and efficient CRISPR-based strategy that enables flexible genome engineering, including the insertion of large tags into the coding region of a gene or the generation of conditional alleles. This strategy should be generally applicable to most *Drosophila* genes. Our results confirmed that about 1 kb homology arms are of sufficient length to insert a large marker cassette, as has been suggested before for other loci^10, 12, 13^. Thus, we could develop an efficient donor plasmid assembly protocol, which facilitates cloning of the donor vector for any gene within a few days. Additionally, our data support the value of a quick pre-testing strategy of predicted sgRNAs in S2 cells to eliminate inefficient sgRNAs, which would likely reduce targeting efficiency *in vivo*. Conveniently, the same *in vitro* transcribed RNAs can be used for both, S2 cell transfections and embryo injections. Our results suggest that indeed a transgenic Cas9 source mediates HDR effectively in *Ligase4* mutant germ line cells. Although *Act5C-Cas9* expression is not restricted to the germ line, injections of the donor vector together with 2 verified sgRNAs led to a targeting efficiency of 2 - 14 % for the incorporation of the large STOP-dsRed cassette, even for the *bent* locus on the heterochromatic 4^th^ chromosome. This suggests that about 50 - 100 fertile G_0_ flies should be sufficient in most cases to identify positive carriers. We thus far deleted up to about 1kb of genomic sequence by HDR. Larger deletions would likely occur at reduced efficiencies, however the dsRed marker should still make it practical to find them. The straight forward identification of carriers together with our simple cloning scheme, this should easily facilitate the insertion of the STOP-dsRed cassette into the gene of choice.

Our 2-step strategy combines the advantages of both, CRISPR and RMCE, and thus allows very flexible modifications of a particular gene region with minimal effort. Multiple fluorescent and affinity tags can be easily inserted or a deleted exon can effectively be replaced by various engineered exon versions. In principle, larger gene parts consisting of multiple exons can also be deleted and replaced by modified versions. This method is particularly valuable for genes that harbour complex transcriptional control and function in many tissues such as *salm* or for genes that are exceptionally large and exhibit complex alternative splicing patterns such as *bent*. The 2-step strategy allows structure-function analysis at the endogenous locus without interfering with the regulatory regions included in introns, which cannot be achieve by simply inserting a cDNA at the transcriptional start site. The functionality of our method was verified by the reversion of the lethality for the step 2 alleles in the first intron of *salm*. This furthermore demonstrates that both steps do not generate additional unintended changes on the chromosome. Therefore, we hope that our strategy will promote the wide application of CRISPR-mediated HDR in *Drosophila*, making it a routine tool used in every fly lab like EMS mutagenesis or P-element mediated transformation were in the last century.

## Methods

### Fly strains and genetics

All fly work, unless otherwise stated, was performed at 25 °C under standard conditions. The *Lig4[169]* null allele^27^ was obtained from the Bloomington *Drosophila* Stock Center, *y[1], M(Act5c-Cas9, [w+])* in *M(3xP3-RFP.attP)ZH-2A*, *w[1118]* was a gift from Fillip Port and Simon Bullock. Both markers (w+ and 3xP3-RFP) were removed by crossing to heat-shock-Cre. The *y[1], M(Act5C-Cas9)ZH-2A, w[1118]* flies were recombined with *Lig4[169]* to obtain *y[1], M(Act5C-Cas9)ZH-2A, w[1118], Lig4[169]*.

### Cell culture

*Drosophila* Schneider 2 (S2) cells stably expressing myc-Cas9 from a ubiquitin promoter were a gift from Klaus Förstemann^19^. S2 cells were cultured in Schneider’s *Drosophila* medium supplemented with 10 % fetal calf serum (Life Technologies) and penicillin/streptomycin (GE Healthcare). sgRNA activities were tested by transfecting 1 µg sgRNA per 24 well into the myc-Cas9 cells using Fugene HD (Promega), followed by DNA extraction and a T7-Endonuclease I assay (see supplied protocol for details).

### Plasmids

CC6-U6-gRNA_hsp70-Cas9 plasmid was a gift from Peter Duchek. pJET1.2-STOP-dsRed: attP1 and splicing acceptor (SA) were amplified with primers XZ82 and XZ83, SV40 terminator with XZ84 and XZ85, and attP2 with XZ88 and XZ89 from DNA extracted from a MiMIC fly line^14^; 3xP3-dsRed was amplified from a fosmid fly line^28^ with primers XZ86 and XZ87. These fragments were cloned into pLR-HD plasmid^29^ by Golden-gate cloning. This assembled attP1-SA-STOP-SV40-3xP3-dsRed-attP2 cassette was amplified with primers XZ195 and XZ196 and blunt cloned into pJET1.2 to generate pJET1.2-STOP-dsRed. As this STOP-dsRed cassette is flanked by two BsmBI sites, it can be easily assembled with both homology arms (Figure 3a): each homology arm of about 1kb was amplified from genomic DNA of the target genotype with Phusion polymerase (NEB) and blunt-end cloned into pJET1.2 (CloneJET PCR Cloning Kit, Thermo Scientific). Primers used to amplify the homology arms have a 5’ BsmBI site enabling Golden Gate assembly with the STOP-dsRed cassette. All primers used are listed in Supplementary Table 1. pBS-donor-backbones: pBS-GGAC-TTCT, pBS-GGAC-ATGC and pBS-CGGA-GTGC were constructed by linearizing pBluescript with KpnI and SacII, followed by amplification with primer pairs XZ150 and XZ151, XZ156 and XZ151, XZ161 and XZ162, respectively, and re-ligation. The generated pBS-donor-backbones harbour two BsmBI sites for donor plasmid assembly. pJET1.2-STOP-dsRed, pJET1.2-HA-left, pJET1.2-HA-right, and an appropriate pBS-backbone were assembled to the pBS-donor vector by Golden-gate cloning. attB plamids: FRT-2xTY1-FRT-V5 and 2xTY1-V5 fragments were synthesized as gBlocks (IDT) and cloned into the attB plasmid for all three reading frames. CC6-U6-gRNA_hsp70-Cas9-sgRNA1,3,4,7,9: the CC6-U6-gRNA_hsp70-Cas9 vector was cut with BbsI (NEB) and the annealed sgRNA targeting oligos were cloned into it. The vas-ΦC31(3xP3-EGFP.attB) plasmid was obtained from Johannes Bischof ^30^. The attB site was removed by digestion with SpeI, followed by re-ligation.

All plasmids for embryo injections were purified with PureLink® HiPure Plasmid Midiprep Kit (Life technologies). Oligos are listed in Supplementary Table 1.

### sgRNA synthesis

The sgRNA dsDNA template was produced using overlap PCR with a small amount of a common sgRNA scaffold primer, a shorter sgRNA amplification primer and a sgRNA gene specific primer that includes the T7 promoter (Figure 3c)^19^. All sgRNA primer sequences are listed in Supplementary Table 1. The PCR product was cleaned by Qiagen MinElute kit (Qiagen). The sgRNAs were transcribed with T7-MEGAshortscript™ Kit (Life Technologies) and purified with MEGAclear™ Transcription Clean-Up Kit (Life Technologies).

### Embryo injection

Pre-blastoderm embryos of the appropriate genotype were de-chorionated and injected with a FemtoJet apparatus (Eppendorf) using self-pulled glass needles (Havard Apparatus) under standard conditions at room temperature. Injected embryos were kept for 2 days at 18 °C and the hatched larvae were collected and grown at 25 °C. For step 1 injections, pBS-donor plasmid, 2 sgRNAs and optionally the CC6-U6-gRNA_hsp70-Cas9 plasmid were mixed and diluted in water. *Lig4[169]* embryos were injected with CC6-U6-gRNA_hsp70-Cas9 plasmid (100 ng / µl) and pBS-donor plasmid (500 ng / µl). *y[1], M(Act5C-Cas9)ZH-2A, w[1118], Lig4[169]* embryos were injected with both sgRNAs (60 - 70 ng / µl, each) and pBS-donor plasmid (500 ng / µl). For step 2 injections the attB plasmid (150 ng/µl) was mixed with vasa-ΦC31 plasmid (200 ng/µl).

**Immunolabelling of IFMs.** Hemi-thoraces of adult *Drosophila* were prepared and stained as described^31^. rabbit anti-Salm was used at 1:50^32^, mouse anti-Flag (Sigma), mouse anti-V5 (Abcam) and rhodamine phalloidin (Invitrogen) were all used at 1:500. Nuclei were visualized by embedding in Vectashield plus DAPI (Vector Laboratories) and images were acquired on a Zeiss LSM780 confocal and processed with FIJI and Photoshop.

### Detailed *Drosophila* genome engineering protocol by CRISPR-RMCE

(Will be deposited at http://www.nature.com/protocolexchange/)

1. **CRISPR-sgRNA design and donor plasmid cloning ∼ 10 days**

1.1 Verify sequence of the targeting region in the fly strain used and in the S2 cells by sequencing to identify potential polymorphisms compared to the published sequence.
1.2 For designing the sgRNA targeting sites, choose one of the web tools^18^. We used an interface designed by the Zhang lab: http://crispr.mit.edu
1.3 sgRNA production

1.3.1 Generate the dsDNA template for sgRNA in vitro transcription as described in^19^.
1.3.2 Transcribe sgRNA by T7-MEGAshortscript™ Kit (AM1354, Life Technologies). Use 150 - 250 ng template for a 20 µl reaction and at 37 °C overnight.
1.3.3 Purify sgRNA by MEGAclear™ Transcription Clean-Up Kit (Life Technologies). Follow the manufacturer’s protocol, and in step 3, add an equal volume of 100 % ethanol to the sample.
1.3.4 Check the sgRNA integrity on a gel and measure the concentration using a photometer (Nanodrop, Thermo Scientific). The expected yield is 100 µg, which is enough for the S2 cell assay and the fly injections.
1.4 sgRNA activity assay in S2-cells

1.4.1 Grow S2 cells in Schneider medium with 10 % FCS (Life Technologies) to 5-10 × 10^6^ cells / ml at 25 °C.
1.4.2 Dilute cells to 0.7 × 10^6^ / ml and plate 1ml cells in S2 medium with 10 % FCS well in a 24 well plate for each transfection.
1.4.3 Prepare the transfection mix by diluting 1 µg sgRNA in 50 µl serum free medium and 4 µl Fugene HD mix plus 46 µl serum free medium, mix both and incubate for 45 min at room temperature.
1.4.4 Add the Fugene/RNA mix to each well and mix gently by pipetting.
1.4.5 After 48 - 60 hours at 25 °C harvest the cells and extract the genomic DNA by QIAamp DNA mini kit (Qiagen).
1.4.6 For the T7 Endonuclease I assay amplify an about 500 bp fragment, which harbours the sgRNA targeting site with Phusion polymerase (NEB), denature and anneal the PCR product as described in^33^.
1.4.7 Mix on ice 10 µl annealed PCR product with 10 µl T7 Endonuclease I master (2 µl T7 endonuclease I buffer, 0.5µl T7 endonuclease I (5 units, NEB) and 7.5 µl water).
1.4.8 Digest at 37 °C for 15 - 20 min using a PCR machine and load on a 1.5 % agarose gel immediately.
1.4.9 Estimate the efficiency of different sgRNAs by comparing the band intensities of the digested and non-digested bands.
1.5 Generation of donor plasmid (can be done in parallel to steps 1.3 and 1.4 to save time)

1.5.1 Amplify left and right homology arms (about 1 kb, start as close to the sgRNA cutting site as possible) with Phusion polymerase (NEB) and clone them into pJET-1.2 according to the CloneJET PCR Cloning Kit (Thermo Scientific).
1.5.2 Assemble the Golden-Gate Cloning reaction (50 ng pBS-backbone, 80 ng pJET1.2-HA-left, 80 ng pJET1.2-HA-right, 80 ng pJET1.2-STOP-dsRed, 1.5 µl 10x T4 ligation buffer, 1 µl BsmBI (NEB, R0580), 1 µl T4 ligase (NEB, M0202) add water to 15 µl).
1.5.3 Ligate in PCR machine using the following cycles: 15 cycles of (37 °C, 5 min and 16 °C, 10 min) / 37 °C, 15 min / 50 °C, 5 min / 80 °C, 5 min / 4°C
1.5.4 Assemble the Plasmid-safe nuclease reaction (15 µl ligation reaction 3 µl 10x Plasmid-safe buffer 1.2 µl 25 mM ATP 1 µl Plasmid-safe nuclease (Epicentre) µl water).
1.5.5 Incubate at 37 °C for 60 min in PCR machine and transform 5 - 10 µl in bacteria. Most growing colonies will be correct.
2. **Fly step 1 - CRISPR-mediated HDR ∼ 6 weeks**

2.1 Inject 600 - 800 *Act5C-Cas9, lig4[169]* embryos with pBS-donor (500 ng / µl) and two sgRNAs (each 60 - 70 ng / µl, targeting close to the chosen homology arms). Collect at least 50 fertile mosaic G_0_ flies.
2.2 Cross G_0_ flies individually (at least 50 vials) either to *yw* flies or appropriate flies and screen all the F_1_ progeny for fluorescent red eyes using a fluorescent binocular (Leica MZ16-FA). Keep track of how of many independent G_0_ founders lead to how many F_1_ carrier flies.
2.3 Generate stocks from an individual F1 carrier by crossing to balancer flies resulting in an isogenised stock for the engineered chromosome. Verify the targeting event by PCR and sequencing.
3. **Fly step 2 - ΦC31-mediated RMCE ∼ 6 weeks**

3.1 Inject a “generic” plasmid generated by^14^ or this study or your own custom-made gene-specific attB plasmid (150 ng / µl) mixed with vasa-ΦC31 plasmid (200 ng / µl) into about 200 embryos from an amplified stock generated at 2.3.
3.2 Cross G_0_ flies individually to an appropriate balancer and screen all F_1_ progeny for non-fluorescent eyes using a fluorescent binocular (Leica MZ16-FA).
3.3 Generate stocks from an individual F1 carrier by crossing to balancer flies resulting in an isogenised stock for the engineered chromosome. Verify the correct orientation of the RMCE by PCR (will be ∼50%).

## Acknowledgements

We thank Klaus Förstemann for the Cas9 expressing S2 cells, Fillip Port and Simon Bullock for the Act5C-Cas9 flies, Peter Duchek for the CC6-U6-gRNA_hsp70-Cas9 plasmid, Johannes Bischof for the ΦC31 plasmid and the Bloomington *Drosophila* Stock Center for fly strains. We are grateful to Reinhard Fässler for generous support and to Bettina Stender, Nicole Plewka and Christiane Barz for excellent technical assistance. We thank Cornelia Schönbauer and Maria Spletter for critical comments on this manuscript. Our work was supported by the Max Planck Society, a Career Development Award from the Human Frontier Science Program (F.S.) and the EMBO Young Investigator Program (F.S.).

---

**Supplementary Table 1.**
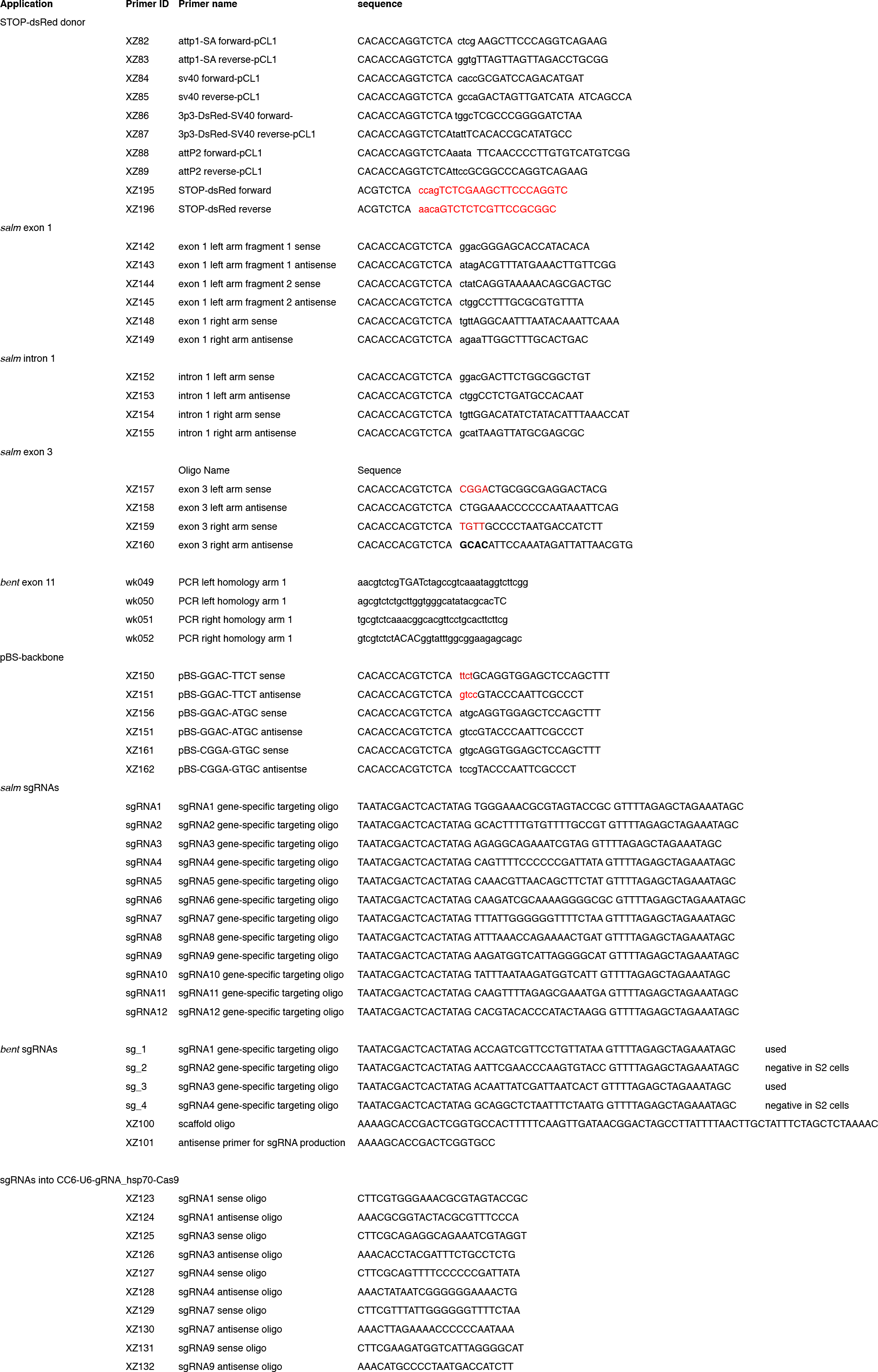
| All primer sequences used in this study.

## References

1. Hsu, P. D., Lander, E. S. & Zhang, F. Development and Applications of CRISPR-Cas9 for Genome Engineering. CELL 157, 1262–1278 (2014).

2. Sander, J. D. & Joung, J. K. CRISPR-Cas systems for editing, regulating and targeting genomes. Nature biotechnology 32, 347–355 (2014).

3. Joung, J. K. & Sander, J. D. TALENs: a widely applicable technology for targeted genome editing. Nature Reviews Molecular Cell Biology 14, 49–55 (2013).

4. Gratz, S. J. et al. Genome engineering of Drosophila with the CRISPR RNA-guided Cas9 nuclease. Genetics 194, 1029–1035 (2013).

5. Sebo, Z. L., Lee, H. B., Peng, Y. & Guo, Y. A simplified and efficient germline-specific CRISPR/Cas9 system for Drosophila genomic engineering. Fly 8, 52–57 (2014).

6. Ren, X. et al. Optimized gene editing technology for Drosophila melanogaster using germ line-specific Cas9. Proceedings of the National Academy of Sciences 110, 19012–19017 (2013).

7. Bassett, A. R., Tibbit, C., Ponting, C. P. & Liu, J.-L. Highly efficient targeted mutagenesis of Drosophila with the CRISPR/Cas9 system. CellReports 4, 220– 228 (2013).

8. Kondo, S. & Ueda, R. Highly improved gene targeting by germline-specific Cas9 expression in Drosophila. Genetics 195, 715–721 (2013).

9. Yu, Z. et al. Highly efficient genome modifications mediated by CRISPR/Cas9 in Drosophila. Genetics 195, 289–291 (2013).

10. Yu, Z. et al. Various applications of TALEN-and CRISPR/Cas9-mediated homologous recombination to modify the Drosophila genome. Biol Open 3, 271–280 (2014).

11. Baena-Lopez, L. A., Alexandre, C., Mitchell, A., Pasakarnis, L. & Vincent, J. P. Accelerated homologous recombination and subsequent genome modification in Drosophila. Development 140, 4818–4825 (2013).

12. Gratz, S. J. et al. Highly Specific and Efficient CRISPR/Cas9-Catalyzed Homology-Directed Repair in Drosophila. Genetics 113.160713 (2014). doi:10.1534/genetics.113.160713/-/DC1

13. Xue, Z. et al. Efficient gene knock-out and knock-in with transgenic Cas9 in Drosophila. G3 (Bethesda) 4, 925–929 (2014).

14. Venken, K. J. T. et al. MiMIC: a highly versatile transposon insertion resource for engineering Drosophila melanogaster genes. Nature Methods 8, 737–743 (2011).

15. Beumer, K. J. et al. Comparing zinc finger nucleases and transcription activator-like effector nucleases for gene targeting in Drosophila. G3 (Bethesda) 3, 1717–1725 (2013).

16. Beumer, K. J., Trautman, J. K., Mukherjee, K. & Carroll, D. Donor DNA Utilization during Gene Targeting with Zinc-finger Nucleases. *G3* (*Bethesda*) (2013). doi:10.1534/g3.112.005439

17. Hadjieconomou, D. et al. Flybow: genetic multicolor cell labeling for neural circuit analysis in Drosophila melanogaster. Nature Methods 8, 260–266 (2011).

18. Beumer, K. J. & Carroll, D. Targeted genome engineering techniques in Drosophila. METHODS 68, 29–37 (2014).

19. Bottcher, R. et al. Efficient chromosomal gene modification with CRISPR/cas9 and PCR-based homologous recombination donors in cultured Drosophila cells. Nucleic Acids Research (2014). doi:10.1093/nar/gku289

20. Schönbauer, C. et al. Spalt mediates an evolutionarily conserved switch to fibrillar muscle fate in insects. Nature 479, 406–409 (2011).

21. Rodriguez-Jato, S., Busturia, A. & Herr, W. Drosophila melanogaster dHCF interacts with both PcG and TrxG epigenetic regulators. PLoS ONE 6, e27479 (2011).

22. Fyrberg, C. C., Labeit, S., Bullard, B., Leonard, K. & Fyrberg, E. Drosophila projectin: relatedness to titin and twitchin and correlation with lethal(4) 102 CDa and bent-dominant mutants. Proc. Biol. Sci. 249, 33–40 (1992).

23. Ayme-Southgate, A., Southgate, R., Saide, J., Benian, G. M. & Pardue, M. L. Both synchronous and asynchronous muscle isoforms of projectin (the Drosophila bent locus product) contain functional kinase domains. Journal of Cell Biology 128, 393–403 (1995).

24. Schnorrer, F. et al. Systematic genetic analysis of muscle morphogenesis and function in Drosophila. Nature 464, 287–291 (2010).

25. Ayme-Southgate, A. & Southgate, R. in *Nature’s Versatile Engine: Insect Flight Muscle Inside and Out* (Vigoreaux, J.) 167–176 (Landes Bioscience), 2006).

26. Gratz, S. J., Wildonger, J., Harrison, M. M. & O’Connor-Giles, K. M. CRISPR/Cas9-mediated genome engineering and the promise of designer flies on demand. Fly 7, 249–255 (2013).

27. McVey, M., Radut, D. & Sekelsky, J. J. End-joining repair of double-strand breaks in Drosophila melanogaster is largely DNA ligase IV independent. Genetics 168, 2067–2076 (2004).

28. Langer, C. C. H., Ejsmont, R. K., Schönbauer, C., Schnorrer, F. & Tomancak, P. In vivo RNAi rescue in Drosophila melanogaster with genomic transgenes from Drosophila pseudoobscura. PLoS ONE 5, e8928 (2010).

29. Cermak, T. et al. Efficient design and assembly of custom TALEN and other TAL effector-based constructs for DNA targeting. Nucleic Acids Research 39, e82–e82 (2011).

30. Bischof, J., Maeda, R., Hediger, M., Karch, F. & Basler, K. An optimized transgenesis system for Drosophila using germ-line-specific φC31 integrases. Proceedings of the National Academy of Sciences 104, 3312 (2007).

31. Weitkunat, M. & Schnorrer, F. A guide to study Drosophila muscle biology. METHODS (2014). doi:10.1016/j.ymeth.2014.02.037

32. Kühnlein, R. P. et al. spalt encodes an evolutionarily conserved zinc finger protein of novel structure which provides homeotic gene function in the head and tail region of the Drosophila embryo. The EMBO Journal 13, 168–179 (1994).

33. Zhang, X., Ferreira, I. R. S. & Schnorrer, F. A simple TALEN-based protocol for efficient genome-editing in Drosophila. METHODS (2014). doi:10.1016/j.ymeth.2014.03.020

